# Pupil size and eye movements differently index effort in both younger and older adults

**DOI:** 10.1101/2024.01.13.575506

**Authors:** Björn Herrmann, Jennifer D. Ryan

## Abstract

The assessment of mental effort is increasingly relevant in neuro-cognitive and lifespan domains. Pupillometry, the measure of the pupil size, is often used to assess effort but has disadvantages. Analysis of eye movements may provide an alternative, but research has been limited to easy and difficult task demands in younger adults. An effort measure must be sensitive to the whole effort profile, including ‘giving up’ effort investment, and capture effort in different age groups. The current study comprised three experiments in which younger (N=66) and older adults (N=44) listened to speech masked by background babble at different signal-to-noise ratios associated with easy, difficult, and impossible speech comprehension. We expected individuals to invest little effort for easy and impossible speech (‘giving up’) but to exert effort for difficult speech. Indeed, pupil size was largest for difficult, but lower for easy and impossible speech. In contrast, gaze dispersion decreased with increasing speech masking in both age groups. Critically, gaze dispersion during difficult speech returned to levels similar to easy speech after sentence offset, when acoustic stimulation was similar across conditions, whereas gaze dispersion during impossible speech continued to be reduced. These findings show that a reduction in eye movements is not a byproduct of acoustic factors, but instead suggest that neuro-cognitive processes, different from arousal-related systems regulating the pupil size, drive reduced eye movements during high task demands. The current data thus show that effort in one sensory domain (audition) differentially impacts distinct functional properties in another sensory domain (vision).

**Significance statement:** The assessment of mental effort is increasingly relevant in many domains. Here, we investigated the sensitivity of a novel measure of effort - the spatial dispersion of eye movements - and compared it to pupillometry, which is a more common effort-assessment tool. Individuals listened to speech masked by background babble at levels associated with easy, difficult, and impossible speech comprehension. Pupil size was largest for difficult, but lower for easy and impossible speech conditions (giving up listening). In contrast, the spatial dispersion of eye movements decreased with increasing speech masking, but this effect was independent of acoustic factors. Instead, the current results suggest that neuro-cognitive processes, different from arousal-related systems regulating the pupil size, drive reduced eye movements during high task demands.

## Introduction

The concept of ‘mental workload’, or effort, is relevant in many neuro-cognitive domains (Brehm & Self, 1989; Hockey, 1997; Kool & Botvinick, 2018). For example, older adults often invest substantial effort to comprehend speech in noise, and this effort is considered an early sign of age-related hearing loss (Eckert, Teubner-Rhodes, & Vaden Jr., 2016; Herrmann & Johnsrude, 2020; Peelle, 2018; Pichora-Fuller et al., 2016). Research is thus increasingly dedicated to quantifying effort (Francis & Love, 2020; Kraus, Tune, Obleser, & Herrmann, 2023; McGarrigle et al., 2014; Moran et al., 2016; Shechter & Share, 2020; Slade, Kramer, Fairclough, & Richter, 2021; A. A. Zekveld, Kramer, & Festen, 2010), but existing effort measures have limitations.

Pupillometry – the measurement of the pupil size – is commonly used to assess effort (van der Wel & van Steenbergen, 2018; M. B. Winn, Wendt, Koelewijn, & Kuchinsky, 2018; Adriana A. Zekveld, Koelewijn, & Kramer, 2018). However, pupil size can vary due to factors unrelated to task demands, such as changes in light (Knapen et al., 2016; Suzuki, Minami, Laeng, & Nakauchi, 2019; Thurman et al., 2021) or in the angle of the eye relative to the eye-tracking camera (Brisson et al., 2013; Fink et al., 2023; Hayes & Petrov, 2016). To address the latter, individuals typically maintain fixation on a point on a computer monitor (Kraus, Obleser, & Herrmann, 2023; Matthew B. Winn & Teece, 2021; Adriana A. Zekveld et al., 2018), but restriction of gaze can reduce memory and mental imagery (Johansson, Holsanova, Dewhurst, & Holmqvist, 2012). Pupillometry has been used to investigate effort in older adults (Nicole D. Ayasse & Wingfield, 2018; Kuchinsky et al., 2013), but aging reduces the dilation range (Guillon et al., 2016; Piquado, Isaacowitz, & Wingfield, 2010; Zhao, Bury, Milne, & Chait, 2019), which could lead to effort misestimations. Moreover, effort-related changes in pupil size are driven by arousal systems in the brain (Joshi & Gold, 2020; Mathôt, 2018; Strauch, Wang, Einhäuser, Van der Stigchel, & Naber, 2022), but arousal can differ between and within populations, adding unwanted variability; for example, arousal can be higher for individuals with hearing loss (Nicolai D. Ayasse & Wingfield, 2020).

Recent work suggests that the quantification of eye movements may provide an alternative effort metric. Eye movements decrease during periods of high compared to low memory load (Kosch, Hassib, Woźniak, Buschek, & Alt, 2018; Walter & Bex, 2021) and with increasing speech masking (Contadini-Wright, Magami, Mehta, & Chait, 2023; Cui & Herrmann, 2023). This research has focused mainly on increases in effort as task demands increase. Yet, disengagement or ‘giving up’ effort investment is common for tasks that feel too effortful, when the reward does not justify the required mental costs (T. S. Braver et al., 2014; Herrmann & Johnsrude, 2020; Westbrook & Braver, 2015). Any measure of effort must be sensitive to disengagement, not only because it is a critical, cognitive transition point (Michael Richter, 2013; Westgate & Wilson, 2018), but also to ensure that the measure is indeed sensitive to variations in effort. The pupil size shows an inverted u-shape: it is largest for difficult listening tasks, but lower during easy and impossible listening (‘giving up’; Ohlenforst et al., 2018; Wendt, Koelewijn, Książek, Kramer, & Lunner, 2018; Adriana A. Zekveld & Kramer, 2014). Whether measures of eye movements also exhibit this u-shaped effort profile associated with easy to difficult to impossible task demands is unclear. Functional differences between pupil and eye movement patterns would suggest distinct neural mechanisms related to effort.

In contrast to measurements of pupil size, measuring reductions in eye movements may be less affected by age-related changes (Bruenech, 2018; Sharpe & Sylvester, 1978; Spooner, Sakala, & Baloh, 1980; Zackon & Sharpe, 1987) as the dynamics related to executing eye movements appear little affected by aging (Abrams, Pratt, & Chasteen, 1998). Further, older adults tend to make more eye movements than younger adults (Liu, Shen, Olsen, & Ryan, 2018; Mazloum-Farzaghi et al., 2022; Ryan, Leung, Turk-Browne, & Hasher, 2007), which may facilitate sensitivity to eye-movement reductions related to effort. Therefore, in the current study, measures of pupil size and the spatial extent of eye movement exploration (i.e., gaze dispersion) were contrasted for the full effort profile in three listening experiments for younger and older adults.

## General methods

### Participants

Younger adults and older adults participated in three experiments. Participants were native English speakers or learned English before the age of 5 years. They reported having normal hearing and did not wear hearing aids or were prescribed one (with the exception of one person whose data were excluded). Participants gave written informed consent prior to the experiment and were paid $7.5 CAD per half-hour for their participation. The study was conducted in accordance with the Declaration of Helsinki, the Canadian Tri-Council Policy Statement on Ethical Conduct for Research Involving Humans (TCPS2-2014), and was approved by the Research Ethics Board of the Rotman Research Institute at Baycrest.

Sample sizes were based on our previous work (Cui & Herrmann, 2023). In Experiment 1, 23 younger adults (median age: 25 years, age range: 18–40 years; 4 male, 17 female, 1 non-binary, 1 trans male) and 20 older adults (median age: 66 years, age range: 56–78 years; 7 male, 13 female) participated. Data from four additional participants were recorded, but not analyzed because they wore hearing aids (n=1), reported having a neurological disease (n=1), their proportion of correct responses for the easiest condition in the behavioral task was equal to or lower than 0.7 (n=1), or more than 50% of trials of their physiological recordings contained more than 50% of missing data (n=1). One person’s pupil data were not included in the analyses, because the person’s pupil data were an extreme outlier (note that inclusion of this data point did not change the significance or absence thereof of statistical tests).

In Experiment 2, 20 younger adults (median age: 23 years, age range: 20–35 years; 10 male, 9 female, 1 non-binary) and 22 older adults (median age: 67 years, age range: 56–77 years; 9 male, 13 female) participated. Data from six additional participants were recorded, but not analyzed because they were a non-native English speaker (n=1), their proportion of correct responses for the easiest condition in the behavioral task was equal to or lower than 0.7 (n=2), or more than 50% of trials of their physiological recordings contained more than 50% of missing data (n=3). One other person indicated having migraines sometimes, but this did not affect their participation. The data of this person were included.

In Experiment 3, 23 younger adults participated (median age: 25 years, age range: 18–33 years; 11 male, 10 female, 2 non-binary). Data from 2 additional participants were recorded but not analyzed because more than 50% of trials contained more than 50% of missing data (n=2). Since younger and older adults showed similar results in the first two experiments, only younger adults were recruited in the last experiment.

### Hearing assessment

Participants sat in a comfortable chair within a sound booth during all experimental procedures. In Experiments 1 and 2, pure-tone audiometric thresholds were obtained for frequencies ranging from 0.25 to 8 kHz using a Maico M28 audiometer. Audiometry was recorded for all older adults, but only for 17 and 9 younger adults for Experiment 1 and 2, respectively, due to misunderstandings related to the execution of the experimental protocols. No audiometry was recorded in Experiment 3, because no older adults were recruited. The pure-tone average threshold was calculated as the mean threshold across 0.5 to 4 kHz frequencies. An independent-samples t-test was used to analyze age-group differences in the pure-tone average threshold. Despite participants self-reporting normal hearing, some adults showed clinically relevant hearing loss (see below). Data from all adults were included nevertheless, because a) our sample is representative of community-dwelling adults who vary in their degree of hearing loss despite self-describing their hearing as normal (Herrmann, Maess, & Johnsrude, 2018, 2023; Moore, 2007; Plack, 2014; Presacco, Simon, & Anderson, 2016), b) they were able to do the behavioral speech-in-noise task, and c) our study seeks to investigate whether eye movements are indicative of listening effort in younger and older adults more generally, rather than investigating differences in people with versus those without hearing loss.

### Experimental setup

Sounds were presented via Sony Dynamic Stereo MDR-7506 headphones and a Steinberg UR22 mkII (Steinberg Media Technologies) external sound card. Experimental procedures were run using Psychtoolbox (v3.0.14) in MATLAB (MathWorks Inc.) on a Lenovo T450s laptop with Microsoft Windows XP. The laptop screen was mirrored to a ViewSonic monitor with a refresh rate of 60 Hz. All sounds were presented at a comfortable listening level that was fixed across participants (∼70-75 dB SPL).

### Stimulus materials and procedure

Stimulus materials consist of sentences from the Harvard sentence lists spoken by a male native English speaker (mean duration: 2.5 s; range of durations: 2–3.1 s) (IEEE, 1969). Speech recordings were taken from a publicly available resource (speaker ID: ‘PNM055’; https://depts.washington.edu/phonlab/projects/uwnu.php; McCloy, Panfili, John, Winn, & Wright, 2018; Panfili, Haywood, McCloy, Souza, & Wright, 2017). Sentences were embedded in a 6.3-second 12-talker babble noise (Bilger, 1984). A sentence started 2.8 s after babble onset (Cui & Herrmann, 2023; Zhang, Malaval, Lehmann, & Deroche, 2022; Figure 1A). For younger adults, sentences were presented at -17 dB (impossible), -4 dB (difficult), and +9 dB SNR (easy). For older adults, SNR levels were increased by 2 dB (−15, -2, +11 dB SNR) relative to younger adults to account for known challenges of older people with the perception of speech in noise (Gustafsson & Arlinger, 1994; Helfer & Freyman, 2008; Herrmann, 2023; Irsik, Johnsrude, & Herrmann, 2022; Presacco, Simon, & Anderson, 2019). We expected listening effort to be low for impossible and easy SNRs, because participants would not invest cognitively for the former and required little effort to comprehend sentences for the latter. Highest listening effort was expected for the difficult SNR. Different SNRs were achieved by adjusting the sound level of the sentence, while keeping the babble level constant across trials (Cui & Herrmann, 2023; Kadem, Herrmann, Rodd, & Johnsrude, 2020; Ohlenforst et al., 2017). This ensured that stimuli with different SNRs did not differ prior to sentence onset (Cui & Herrmann, 2023; Kadem et al., 2020; Ohlenforst et al., 2017). After each stimulus, a probe word occurred on the screen. Participants were asked to indicate whether the probe word was semantically related or unrelated to the sentence (Cui & Herrmann, 2023; Kadem et al., 2020; Rodd, Johnsrude, & Davis, 2010; Rodd, Longe, Randall, & Tyler, 2010).

**Figure 1:**
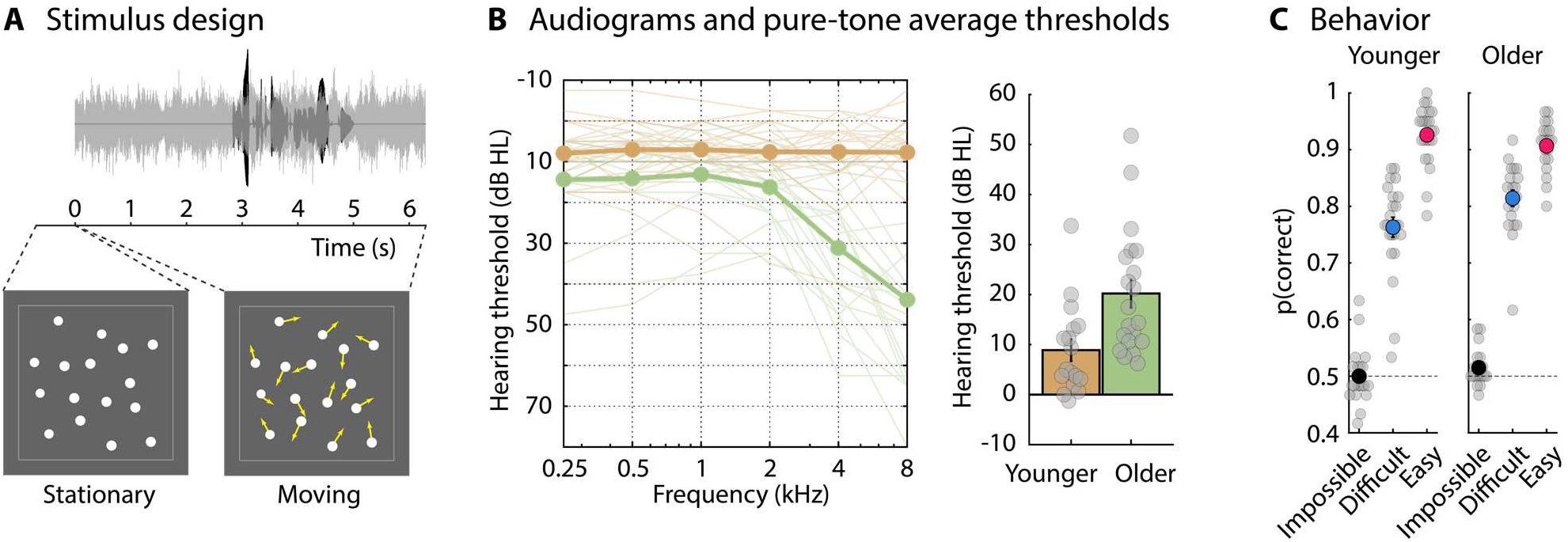
Experimental design, pure-tone audiometric thresholds, and behavioral data. A: Stimulation paradigm. A sentence was presented in a 6.3-s babble noise. The sentence onset was at 2.8 s and the averaged sentence offset was at 5.3 s. A moving-dot display was presented while participants listened to the auditory stimulus. Following a 0.7-s stationary display prior to babble onset, sixteen dots smoothly moved on the screen throughout the entire duration of the auditory stimulus. No task was required using the moving-dot display. B: Left: Pure-tone audiometric thresholds: The thin lines reflect data from each individual participant. Thick lines reflect the mean across participants. Right: Pure-tone average hearing threshold (mean across 0.5, 1, 2, and 4 kHz). Gray dots reflect the pure-tone average threshold for individual participants. Audiograms were available for 17 out of 23 younger and for 20 out of 20 older adults. C: Proportion of correct responses in the semantic-relatedness task for younger (left) and older adults (right). Gray dots reflect data from individual participants. The dashed horizontal line at 0.5 reflects chance level. Error bars reflect that standard error of the mean.

Participants listened to 60 sentences per SNR condition across 5 experimental blocks. Within each block, participants listened to 12 sentences per SNR condition. The order of SNR conditions was randomized such that each SNR level occurred maximally two times in direct succession (Experiment 1 and 2) or was blocked with 12 sentences of the same SNR in direct succession (Experiment 3). Assignment of SNR levels to specific sentences was randomized across participants to avoid confounding specific sentences with SNR levels. Participants underwent a short 12-trials training block to familiarize them with the eye-tracking calibration procedures and the semantic-relatedness task. Different sentences from the Harvard sentence lists were used in Experiments 1, 2, and 3.

Pupillometry studies typically require a participant to fixate centrally on a fixation point on a computer screen to reduce eye movements (Farahani et al., 2020; Kadem et al., 2020; Matthew B. Winn & Teece, 2021; Adriana A. Zekveld, Heslenfeld, Johnsrude, Versfeld, & Kramer, 2014; Adriana A. Zekveld et al., 2018; Zhao et al., 2019). However, an investigation into the degree to which eye movements reflect listening effort may be impeded by artificially restricting eye movements to a fixation point (Cui & Herrmann, 2023), and the restriction of eye movements can reduce behavioral speech-comprehension effects (Cui & Herrmann, 2023). To facilitate eye movements, similar to our recent work, we presented an incidental multi-object movement display (Cui & Herrmann, 2023; see other work using object tracking, Alvarez & Franconeri, 2007; Cavanagh & Alvarez, 2005; Herrmann & Johnsrude, 2018a, 2018b; Scholl, 2009). To this end, 16 white dots [dot diameter: 1.2 cm (0.9°)] were presented on gray background and moved on the screen during the presentation of the auditory stimuli. Presentation of dots was constrained to a display frame of 20.6 cm width (15.6°) and 19.4 cm height (14.7°) centered on the screen and highlighted to the participants by a light gray frame (Figure 1A). Dots never moved outside of the display frame and never overlapped during movements; dots moved ∼3.7 cm/s (2.8°/s). The dot display occurred 0.7 s prior to babble onset and remained stationary for the 0.7 s duration. Dot movements started with the onset of the babble and continued throughout the 6.3 s duration of the auditory stimulus, at which point the dots disappeared. Critically, participants were instructed that their task was to comprehend the sentence and perform the semantic-relatedness task. No task was required for the dot display. Participants were instead instructed to look at the screen in whatever way they wanted (Cui & Herrmann, 2023; Johansson et al., 2012; Johansson, Holsanova, & Holmqvist, 2006; Johansson, Holsanova, & Homqvist, 2011).

The three experiments of the current study differed in the degree to which individuals could anticipate the comprehension difficulty of an upcoming sentence. In Experiment 1, no cues were given to participants, and they therefore only knew about the difficulty of a sentence upon sentence onset. Importantly, anticipation of the level of difficulty is a critical component in effort frameworks to help an individual to know when to disengage to preserve effort and reduce fatigue (Brehm & Self, 1989; Herrmann & Johnsrude, 2020; Michael Richter, 2013; M. Richter, Gendolla, & Wright, 2016). In Experiment 2, a visual cue indicated the difficulty of an upcoming sentence. To this end, the 16 dots of the 0.7-s stationary dot-display were color-coded, with green dots indicating easy comprehension, yellow dots indicating difficult comprehension, and red dots indicating impossible comprehension. Upon babble onset, the color of the dots turned white and all dots started to move. The color to difficulty assignment was not counter-balanced across participants, because we wanted to avoid confusing color-difficulty associations (i.e., red indicating easy) and we reasoned the cue color is unlikely to influence ocular responses 2.8 s later (sentence onset). We anticipated that color cues would enable participants to regulate effort investment on a sentence-by-sentence basis. In Experiment 3, SNR conditions (easy, difficult, impossible) were presented in mini-blocks of 12 sentences during an experimental block. Prior to each mini-block, participants were also visually presented with the text “Comprehension of the next few sentences will likely be: …” completed by either “easy”, “difficult”, or “impossible” depending on the SNR condition. The dots were white throughout and did not cue SNR conditions. By blocking SNR conditions, we reasoned that participants would more easily be able to regulate effort investment than in Experiment 2.

### Behavioral data analysis

The proportion of correct responses in the semantic-relatedness task was calculated for each SNR condition. Behavioral and physiological data from participants who had a proportion of correct responses equal to or lower than 0.7 in the easiest condition were excluded, based on the assumption that, in such cases, participants either did not pay sufficient attention or experienced undue difficulties. A repeated measures analysis of variance (rmANOVA) was calculated using the proportion of correct responses as dependent measure. SNR (easy, difficult, impossible) was included as a within-participants factor and age group (younger, older) was included as a between-participants factor (with the exception of Experiment 3, which only included younger adults). Post hoc tests were calculated for significant main effects and interactions. Holm’s methods was used to correct for multiple comparisons (Holm, 1979).

### Pupillometry and eye-movement recordings

During the experiments, participants placed their head on a chin and forehead rest facing the computer monitor at a distance of about 70 cm. Pupil area and eye movements were recorded continuously from the right eye (or the left eye if the right eye could not be tracked accurately) using an integrated infrared camera (EyeLink 1000 eye tracker; SR Research Ltd.) at a sampling rate of 500 Hz. Nine-point fixation was used for eye-tracker calibration prior to each block (McIntire, McIntire, McKinley, & Goodyear, 2014).

### Processing of pupil-size and eye-movement data

The main metrics of the current study were pupil size and gaze dispersion (Cui & Herrmann, 2023). Preprocessing was calculated for each block of presentation separately. Preprocessing of pupil-size and eye-movement data involved removing eye blinks and other artifacts. For each eye blink indicated by the eye tracker, all data points of the pupil time course and the x and y eye-coordinate time courses between 100 ms before and 200 ms after a blink were set to NaN (‘not a number’ in MATLAB). In addition, pupil-size values that differed from the mean pupil size (across a block) by more than 3 times the standard deviation were classified as outliers and the corresponding data points of the pupil time course and the x and y eye-coordinate time courses were set to NaN. Missing pupil data (coded as NaN) resulting from artifact rejections and outlier removal were interpolated using MATLAB’s ‘pchip’ method. Pupil-size time courses were filtered with a 5-Hz low-pass filter (51 points, Kaiser window; β = 4). X and y eye-coordinate time courses were not interpolated. That is, missing data points (NaNs) were ignored in the calculation of the eye-movement metric (i.e., gaze dispersion).

Changes in eye movements were investigated using spatial gaze dispersion (Cui & Herrmann, 2023). Spatial gaze dispersion is a measure of the general tendency for the eyes to move around. It was calculated as the standard deviation in gaze across time points, averaged across x- and y-coordinates, and transformed to logarithmic values. Smaller values indicate less gaze dispersion. To obtain time courses for gaze dispersion, it was calculated for 1-s sliding time windows centered sequentially on each time point. If more than 90% of data were unavailable within a 1-s time window (that is, 450 or more samples were NaN-coded), gaze dispersion for the corresponding time point was set to NaN and ignored during averaging (Cui & Herrmann, 2023). Gaze dispersion is a broad measure that captures any eye movements. Analyses of saccades (including microsaccades) require setting a velocity threshold to determine what counts as a saccade (Contadini-Wright et al., 2023; Cui & Herrmann, 2023; Engbert, 2006; Engbert & Kliegl, 2003; Kadem et al., 2020; Widmann, Engbert, & Schröger, 2014), but the threshold can influence whether or not effects of cognition on saccades can be observed (Cui & Herrmann, 2023). Nonetheless, we explored the sensitivity of saccades to changes in SNR and results were very similar to the results for gaze dispersion.

Continuous pupil-size and gaze-dispersion data were divided into single-trial time courses ranging from -1.7 to 6.3 s time-locked to babble onset. Data for an entire trial were excluded from analysis if the percentage of NaN data entries made up more than 50% of the trial. Pupil size and gaze dispersion were averaged across trials, separately for each SNR condition. Pupil size was also normalized relative to the -1.7 s to -0.7 s baseline period (i.e., the period prior to the onset of the visual dot display), by subtracting the mean baseline pupil size from the pupil size at each time point. Gaze dispersion does not require baseline correction.

For statistical analyses, gaze dispersion was averaged across the 2.8–5.3-s time window during which sentences were presented (sentences started at 2.8 s after babble onset; average sentence offset was at 5.3 s). Mean pupil size was calculated for a time window that was delayed by 0.5 s to 3.3–5.8-s, because the pupil size is known to change relatively slowly and peak late during sentence listening (Kadem et al., 2020; Knapen et al., 2016; Matthew B. Winn & Moore, 2018; M. B. Winn et al., 2018; Zhang et al., 2022). We also analyzed data after sentence offset, focusing on the 5.3 s to 6 s time window, to investigate whether gaze dispersion is sensitive to decreased listening effort when speech has ended (we ignored the last 0.3 s prior to babble offset to avoid impacting the analyses by back smearing of eye movement activity following babble offset). Separate rmANOVAs were calculated for different measures (pupil size, gaze dispersion) and time windows, using the within-participants factor SNR (easy, difficult, impossible) and the between-participants factor Age Group (younger, older). Age Group was not included in Experiment 3, because only younger adults were recruited. Post hoc tests were calculated for significant main effects and interactions. Holm’s methods was used to correct for multiple comparisons (Holm, 1979). We also examined linear and quadratic SNR effects. A quadratic effect in the absence of a linear effect would be consistent with the expected inverted u-shaped effort profile. Statistical analyses were carried out in JASP software (v0.18.1; JASP, 2023). Note that JASP uses pooled error terms and degrees of freedom from an rmANOVA for the corresponding post hoc, linear, and quadratics effects. The reported degrees of freedom are thus higher than for direct contrasts had they been calculated independently from the rmANOVA.

## Results for Experiment 1

Younger and older adults participated in Experiment 1. Pure-tone audiometric thresholds were higher in older compared to younger adults (t_35_ = 3.099, p = 0.004, d = 1.022), as expected, and some adults showed clinically relevant hearing loss (Figure 1B). Data from these individuals were included in the analyses nonetheless as described above.

Participants listened to sentences in background babble at three different SNR levels that were selected to make sentence comprehension easy, difficult, or impossible. No prior cues were provided to participants about whether comprehension for an upcoming sentence would be easy, difficult, or impossible. Participants judged whether the probe word that followed a sentence was semantically related to the sentence. Performance in the semantic relatedness task was higher for easy than difficult (t_82_ = 10.974, p_Holm_ = 9.1 · 10^-18^, d = 2.248), easy than impossible (t_82_ = 35.119, p_Holm_ = 1 · 10^-50^, d = 7.193), and difficult than impossible SNRs (t_82_ = 24.144, p_Holm_ = 1 · 10^-38^, d = 4.945; effect of SNR: F_2,82_ = 645.578, p = 6.6 · 10^-51^, ω^2^ = 0.902), as expected. Performance was higher in older than younger adults for the difficult condition (t_82_ = 2.942, p_Holm_ = 0.012, d = 0.899; SNR × Age Group interaction: F_2,82_ = 4.590, p = 0.013, ω^2^ = 0.049), but not for the other SNR conditions (for all p_Holm_ > 0.5). Differences between SNR conditions were significant for both age groups (for all p_Holm_ < 0.001). There was no effect of Age Group (p = 0.165). Hence, the SNR manipulation influenced behavior as intended, but older adults performed better than younger adults on the difficult condition (Figure 1C).

The pupil size was larger during the difficult than the impossible SNR (t_80_ = 3.107, p_Holm_ = 0.008, d = 0.276) and numerically, but non-significantly larger for the difficult than the easy SNR (t_80_ = 1.981, p_Holm_ = 0.102, d = 0.176), whereas easy and impossible conditions did not differ (p_Holm_ = 0.264; effect of SNR: F_2,80_ = 4.947, p = 0.009, ω^2^ = 0.011; Figure 2). The quadratic (t_80_ = -2.937, p = 0.004) but not the linear (p = 0.264) SNR effect was also significant. This quadratic SNR effect is the expected inverted u-shaped profile that indicates sensitivity of the pupil size to listening effort (cf. Ohlenforst et al., 2018; Ohlenforst et al., 2017; Wendt et al., 2018; Adriana A. Zekveld & Kramer, 2014). There was also a transient increase in the pupil size for the easy SNR following sentence onset (Figure 2A). This transient increase may have been due to increased arousal following the unpredictable sound-level increase associated with the high SNR of the sentence (+9 dB SNR and +11 dB SNR for younger and older adults, respectively), and may have made observing a difference between the difficult and easy SNRs more challenging. There was no effect of Age Group (F_1,40_ = 3.010, p = 0.090, ω^2^ = 0.024) and no Age Group × SNR interaction (F_2,80_ = 0.060, p = 0.942, ω^2^ < 0.001).

**Figure 2:**
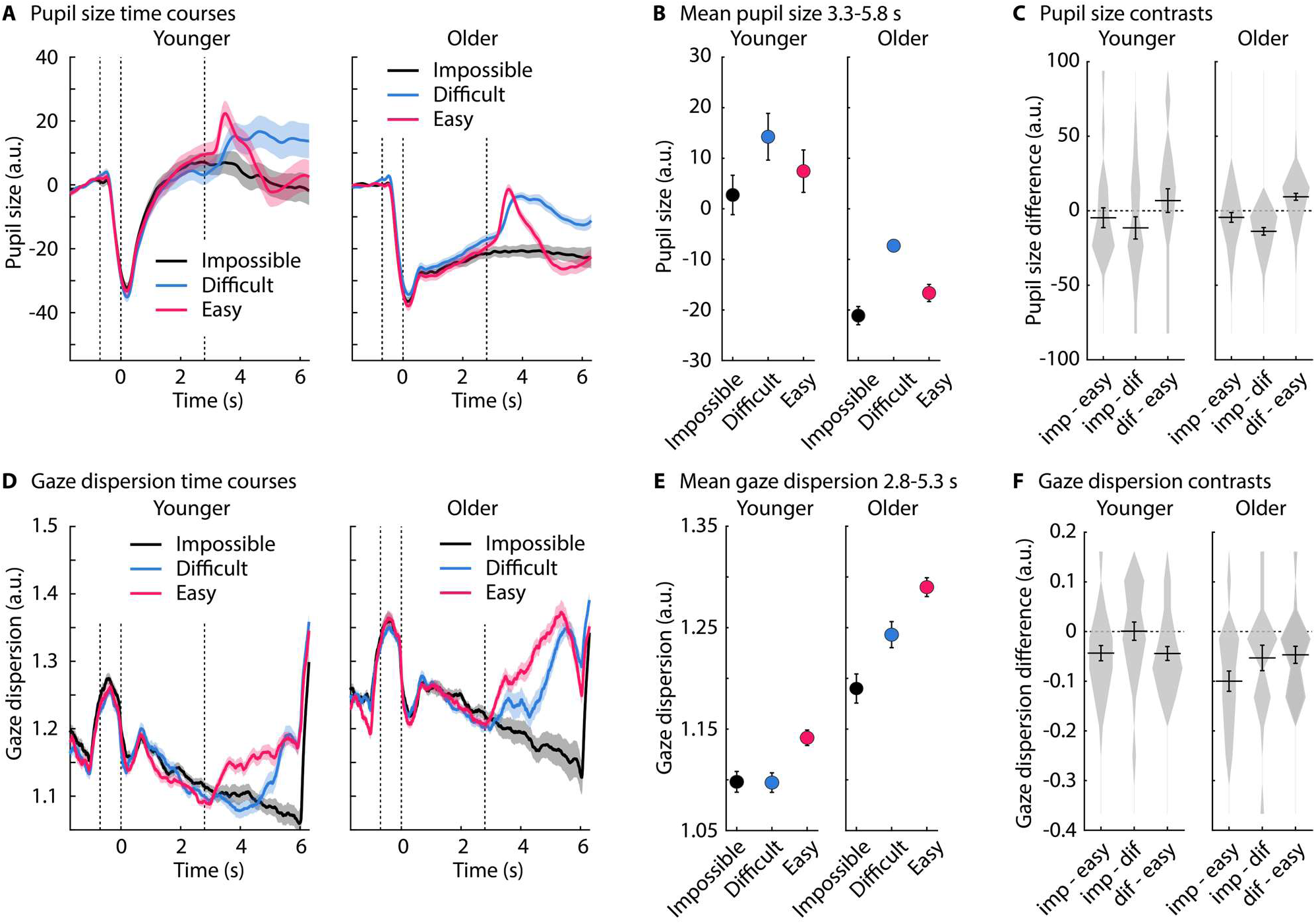
Pupil size and gaze dispersion for Experiment 1. A: Pupil-size time courses. The shaded area reflects the standard error of the mean (between-participant variance removed; Masson & Loftus, 2003). The dashed vertical lines indicate, from left to right, the onset of the dot display (stationary), onset of the babble noise and dot movements, sentence onset. B: Mean pupil size across the 3.3–5.8 s time window (sentence onset at 2.8 s and average sentence offset at 5.3 s). Error bars reflect the standard error of the mean (removal of between-participant variance; Masson & Loftus, 2003). C: Difference in pupil size between different SNR conditions. Violin plots reflect the histogram of individual data points. Horizontal lines reflect the mean. Error bars reflect the standard error of the mean. D-F: The same types of data plots as in panels A-C are depicted for gaze dispersion. Panel E and F reflect mean gaze dispersion across the 2.8-5.3 s time window.

Gaze dispersion was also sensitive to SNR (F_2,82_ = 15.240, p = 2.4 · 10^-6^, ω^2^ = 0.016), but differently than the pupil size. Gaze dispersion decreased when speech comprehension was difficult (t_82_ = 3.465, p_Holm_ = 0.002, d = 0.204) and impossible (t_82_ = 5.455, p_Holm_ = 1.5 · 10^-6^, d = 0.321) relative to when comprehension was easy. Eye movements were further reduced under the impossible compared to the difficult condition (t_82_ = 1.99, p_Holm_ = 0.05, d = 0.117). This was also reflected in the linear (t_82_ = 5.455, p = 5.7 · 10^-11^) SNR effect; there was no quadratic effect (t_82_ = 0.851, p = 0.397). There was no effect of Age Group (F_1,41_ = 3.733, p = 0.060, ω^2^ = 0.032) nor an Age Group × SNR interaction (F_2,82_ = 2.942, p = 0.058, ω^2^ = 0.002). Critically, the reduction in eye movements associated with SNR conditions is not related to SNR per se. That is, eye movements remained reduced for the impossible relative to the difficult SNR (t_82_ = 7.601, p_Holm_ = 1.3 · 10^-10^, d = 0.751) and the easy SNR (t_82_ = 7.537, p_Holm_ = 1.3 · 10^-10^, d = 0.745) even after a sentence ended (5.3 to 6 s) and no acoustic differences between the three SNR conditions were present, whereas eye movements did not differ between the easy and the difficult conditions after sentence offset (t_82_ = 0.064, p_Holm_ = 0.949, d = 0.006; effect of SNR: F_2,82_ = 38.193, p = 1.9 × 10^-12^, ω^2^ = 0.110). This suggests that a cognitive process, rather than acoustic differences, led to reduced eye movements for the impossible condition and the dissociation from the pattern of results for the pupil size.

The results of Experiment 1 show that pupil size indexes listening effort as expected. That is, pupil size increased for difficult speech comprehension, but not when comprehension was impossible (Ohlenforst et al., 2018; Ohlenforst et al., 2017; Wendt et al., 2018; Adriana A. Zekveld & Kramer, 2014). In contrast, eye movements, as indexed by gaze dispersion, decreased for both, when comprehension was difficult and when it was impossible, suggesting different processes underlie changes in pupil size and eye movements during listening. In Experiment 1, participants were unable to anticipate whether comprehension of an upcoming sentence will be easy, difficult, or impossible. However, anticipating the difficulty of a task is a critical component in frameworks that are concerned with motivation and cognitive control (Brehm & Self, 1989; Michael Richter, 2013; M. Richter et al., 2016). Anticipating when listening is challenging can help regulate how much effort is invested. In Experiment 1, listeners may have continued attentive listening even under impossible conditions, affecting their eye movements, but not arousal-related changes in pupil size, because they were unable to predict ahead of time whether a sentence would be understood. Experiment 2 was designed to investigate whether prior knowledge about the comprehension difficulty changes the influence of SNR on pupil responses and eye movements.

## Experiment 2

Younger and older adults participated in Experiment 2. Pure-tone audiometric thresholds were higher in older compared to younger adults (t_29_ = 4.04, p = 3.6 · 10^-4^, d = 1.599; Figure 3A,B), as expected. Participants listened to sentences in background babble at three different SNR levels that made comprehension easy, difficult, or impossible. Critically, in Experiment 2, a visual cue prior to babble onset indicated whether comprehension of the upcoming sentence would be easy, difficult, or impossible. As for Experiment 1, performance decreased with SNR (effect of SNR: F_2,80_ = 646.899, p = 4 · 10^-50^, ω^2^ = 0.908). Older adults exhibited higher performance than younger adults for the difficult SNR (t_80_ = 2.899, p_Holm_ = 0.013, d = 0.896), but not the easy or impossible SNRs (for both p_Holm_ > 0.2; Age Group × SNR interaction: F_2,80_ = 5.719, p = 0.005, ω^2^ = 0.067; Figure 3C). There was no effect of Age Group (p = 0.607).

**Figure 3:**
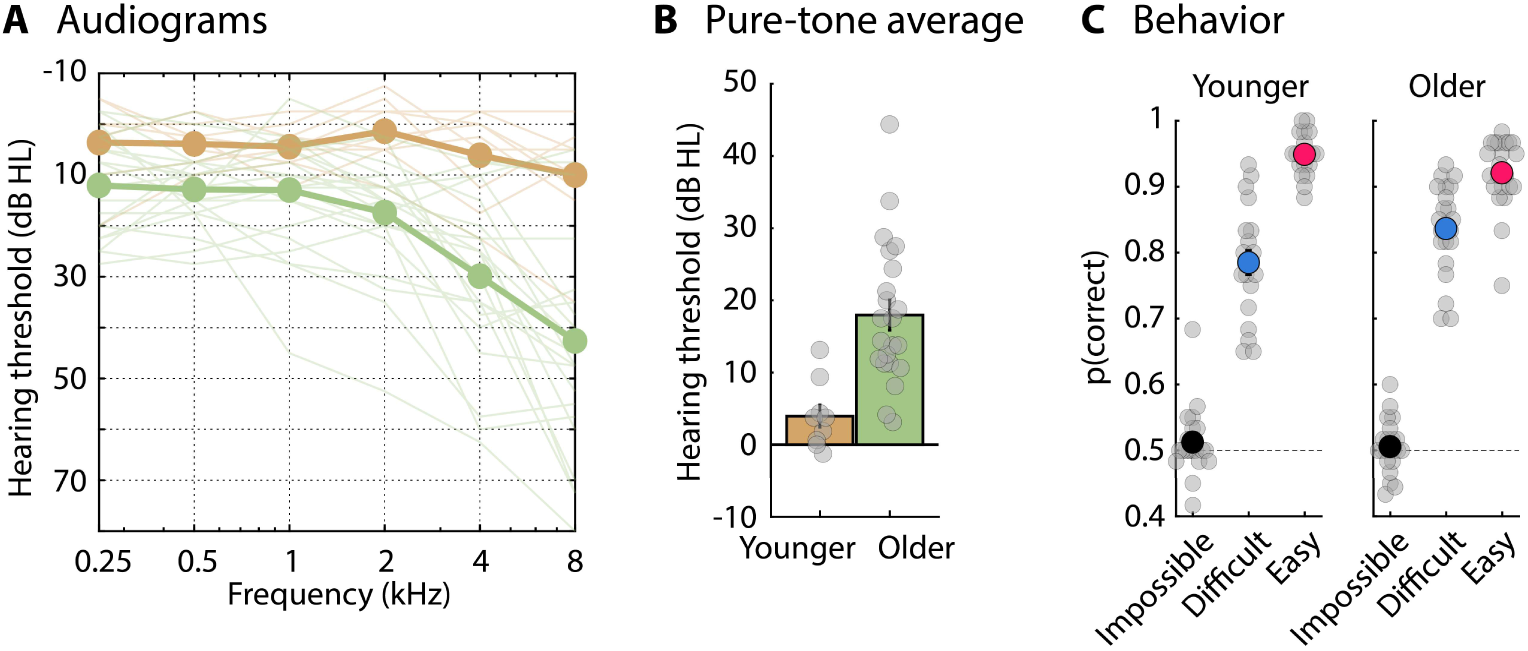
Pure-tone audiometric thresholds and behavioral data for Experiment 2. A: Pure-tone audiometric thresholds: The thin lines reflect data from individual participants. Thick lines reflect the mean across participants. Note that in Experiment 2, audiograms were recorded for all older adults, but only for 9 younger adults. B: Pure-tone average hearing threshold (mean across 0.5, 1, 2, and 4 kHz). Gray dots reflect the pure-tone average threshold for individual participants. C: Proportion of correct responses in the semantic-relatedness task for younger (left) and older adults (right). Gray dots reflect data from individual participants. The dashed horizontal line at 0.5 reflects chance level. Error bars reflect that standard error of the mean.

Pupil size was modulated by SNR (F_2,80_ = 8.542, p = 4.3 · 10^-4^, ω^2^ = 0.037), such that pupil size was larger under difficult than easy (t_80_ = 2.918, p_Holm_ = 0.009, d = 0.359) and impossible conditions (t_80_ = 3.994, p_Holm_ = 4.3 · 10^-4^, d = 0.491), whereas the pupil size for easy and impossible SNRs did not differ (t_80_ = 1.077, p_Holm_ = 0.285, d = 0.132). The quadratic (t_80_ = -3.991, p = 1.5 · 10^-4^) but not the linear (p = 0.285) SNR effect was also significant. Hence, pupillometry showed the expected inverted u-shaped profile, indicating sensitivity to listening effort (Figure 4; Ohlenforst et al., 2018; Ohlenforst et al., 2017; Wendt et al., 2018; Adriana A. Zekveld & Kramer, 2014). There was no age-group difference no an interaction involving age group (ps > 0.5).

**Figure 4:**
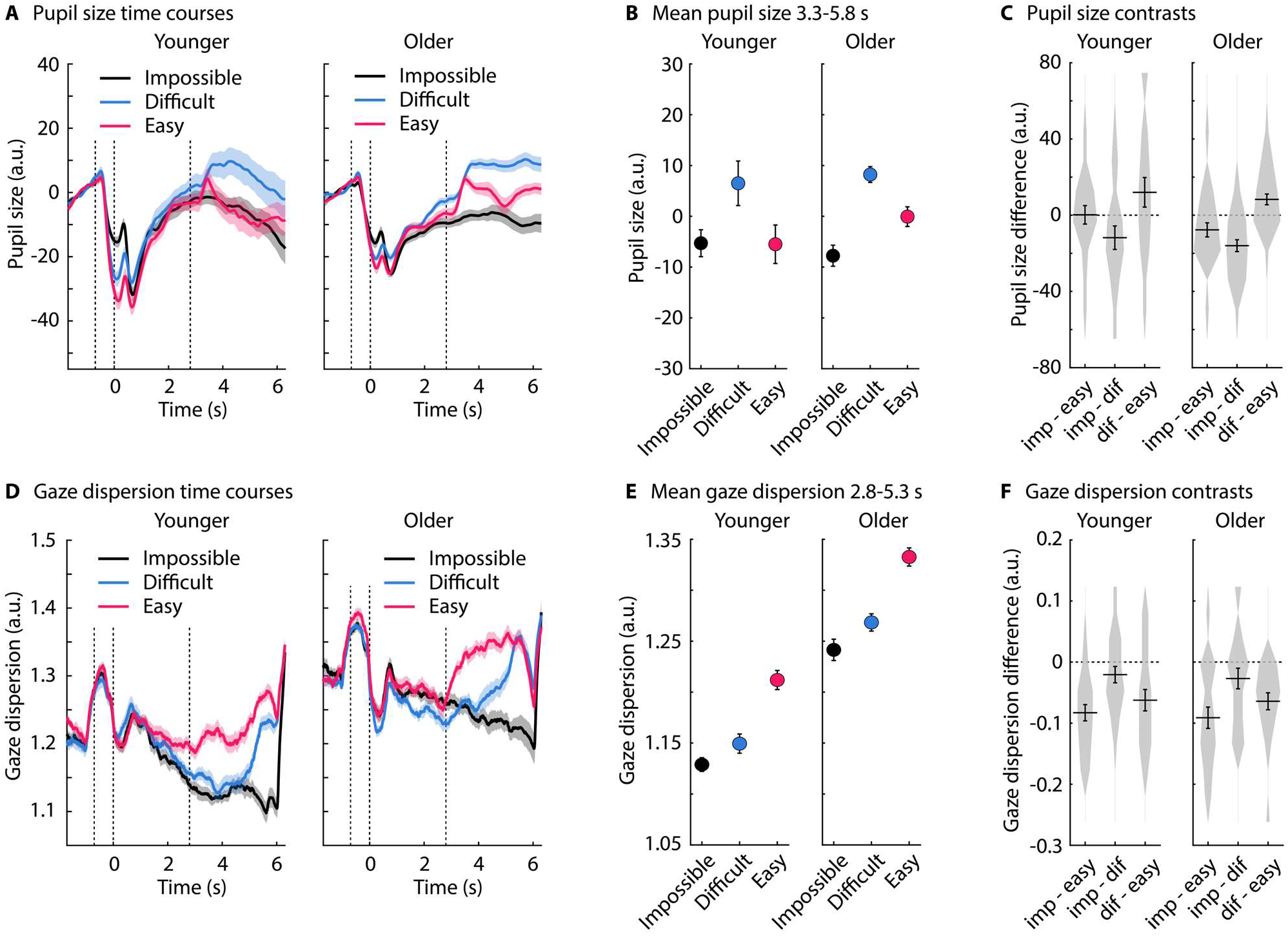
Pupil size and gaze dispersion for Experiment 2. A: Pupil size time courses. The shaded area reflects the standard error of the mean (between-participant variance removed; Masson & Loftus, 2003). The dashed vertical lines indicate, from left to right, the onset of the dot display (stationary), onset of the babble noise and dot movements, sentence onset. B: Mean pupil size across the 3.3–5.8 s time window (sentence onset at 2.8 s and average sentence offset at 5.3 s). Error bars reflect the standard error of the mean (removal of between-participant variance; Masson & Loftus, 2003). C: Difference in pupil size between different SNR conditions. Violin plots reflect the histogram of individual data points. Horizontal lines reflect the mean. Error bars reflect the standard error of the mean. D-F: The same types of data plots as in panels A-C are depicted for gaze dispersion. Panel E and F reflect mean gaze dispersion across the 2.8-5.3 s time window.

Gaze dispersion also showed the same pattern as in Experiment 1. During sentence presentation (2.8 to 5.3 s), gaze dispersion decreased with decreasing SNR (F_2,80_ = 33.267, p = 3 · 10^-11^, ω^2^ = 0.030), and there was no age-group difference or interaction (ps > 0.05). Specifically, gaze dispersion was reduced for difficult (t_80_ = 5.738, p_Holm_ = 3.3 · 10^-7^, d = 0.303) and impossible conditions (t_80_ = 7.890, p_Holm_ = 4 · 10^-11^, d = 0.416) compared to the easy condition, and gaze dispersion was further reduced for the impossible compared to the difficult condition (t_80_ = 2.151, p_Holm_ = 0.034, d = 0.114). This was also reflected in the linear (t_80_ = 7.890, p = 1.3 · 10^-11^) and quadratic (t_80_ = 2.071, p = 0.042) SNR effects. Moreover, gaze dispersion remained lower when comprehension was impossible, compared to both the easy SNR (t_80_ = 8.218, p_Holm_ = 9.1 · 10^-12^, d = 0.690) and the difficult SNR (t_80_ = 7.298, p_Holm_ = 3.8 · 10^-10^, d = 0.612), even after sentence offset (5.3–6 s) when no acoustic differences between conditions were present, whereas eye movements did not differ between easy and difficult SNRs after sentence offset (p_Holm_ = 0.36; effect of SNR: F_2,80_ = 40.545, p = 6.9 · 10^-13^, ω^2^ = 0.087). This suggests, similar to Experiment 1, that cognitive processes, rather than acoustic differences, underlie the pattern of eye movements.

In Experiment 2, participants could anticipate the level of speech comprehension difficulty prior to sentence onset, but SNRs associated with easy, difficult, and impossible comprehension randomly varied from trial to trial. This trial-to-trial change in SNR may reduce individuals’ ability to fully make use of anticipatory processes. In some previous pupillometry work that investigated the relation between SNR and pupil size, trials with different SNRs were presented in separate blocks (Wendt et al., 2018). This may facilitate the anticipation of upcoming comprehension difficulties and may better enable individuals to give up listening (impossible) or ready themselves for sustained effort (difficult). Experiment 3 was designed to test whether a blocked presentation of SNRs changes the influence of SNR on eye movements.

## Experiment 3

In Experiment 3, younger adults listened to sentences in background babble at three different SNR levels that made comprehension easy, difficult, or impossible. SNR conditions were blocked such that 12 trials of the same SNR condition occurred in direct succession. Each of the 12-trial blocked presentations was preceded by written information about the comprehension difficulty of the upcoming sentences. Performance in the semantic-relatedness task decreased with decreasing SNR (F_2,44_ = 272.306, p = 1.7 · 10^-25^, ω^2^ = 0.865; all post hoc contrasts p_Holm_ < 0.001; Figure 5A).

**Figure 5:**
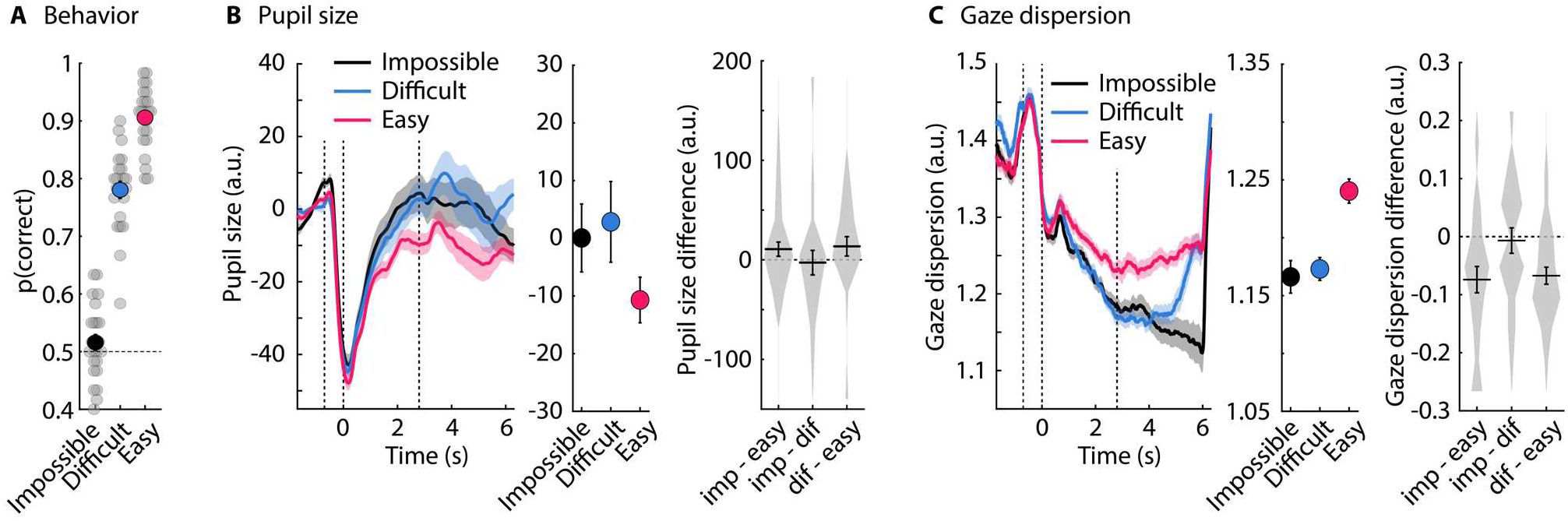
Behavioral data, pupil area, and gaze dispersion for Experiment 3. A: Proportion of correct responses in the semantic-relatedness task for younger adults. Gray dots reflect data from individual participants. The dashed horizontal line at 0.5 reflects chance level. B: Pupil size data. Left: time courses. The shaded area reflects the standard error of the mean (between-participant variance removed; Masson & Loftus, 2003). The dashed vertical lines indicate, from left to right, the onset of the dot display (stationary), onset of the babble noise and dot movements, sentence onset. Middle: Mean pupil size across the 3.3–5.8 s time window (sentence onset at 2.8 s and average sentence offset at 5.3 s). Error bars reflect the standard error of the mean (removal of between-participant variance; Masson & Loftus, 2003). Right: Difference in pupil size between different SNR conditions. Violin plots reflect the histogram of individual data points. Horizontal lines reflect the mean. Error bars reflect the standard error of the mean. C: The same data plots as in panel B are depicted for gaze dispersion (mean across the 2.8-5.3 s time window).

There was no effect of SNR on pupil size (F_2,44_ = 1.032, p = 0.365, ω^2^ < 0.001; Figure 5B), nor a linear or quadratic SNR effect (both ps > 0.25). In contrast, gaze dispersion was smaller for the difficult (t_44_ = 3.352, p_Holm_ = 0.003, d = 0.313) and the impossible SNR (t_44_ = 3.692, p_Holm_ = 0.002, d = 0.345) compared to the easy SNR, whereas the difficult and the impossible SNRs did not differ (p_Holm_ > 0.7; effect of SNR: F_2,44_ = 8.327, p = 8.6 · 10^-4^, ω^2^ = 0.021; Figure 5C). This was also reflected in the linear SNR effect (t_44_ = 3.692, p = 6.1 · 10^-4^; quadratic: t_44_ = 1.739; p = 0.089). As for Experiments 1 and 2, the reduction in gaze dispersion cannot be due to acoustic differences, because gaze dispersion remained reduced for the impossible SNR compared to both the easy (t_44_ = 3.939, p_Holm_ = 8.7 · 10^-4^, d = 0.610) and the difficult SNR (t_44_ = 3.481, p_Holm_ = 0.002, d = 0.539) even after sentence offset (5.3–6 s; effect of SNR: F_2,44_ = 9.281, p = 4.3 · 10^-4^, ω^2^ = 0.062). Gaze dispersion did not differ between the difficult and the easy SNR conditions after sentence offset (p_Holm_ > 0.6).

Interestingly, Figure 5C also indicates that gaze dispersion was already reduced for both the difficult and impossible SNRs, compared to the easy SNR, prior to sentence onset, likely because blocking SNR conditions enabled participants to anticipate the degree of listening challenge of the upcoming sentence. Indeed, in the 1.8 to 2.3 s time window (sentence onset at 2.8 s; 2.3–2.8 s was excluded to avoid smearing post-sentence onset effects back due to the sliding window calculation of gaze dispersion), the SNR effect was significant (F_2,44_ = 3.382, p = 0.043, ω^2^ = 0.005), and eye movements were reduced for both the difficult and impossible SNRs compared to the easy SNR, although only marginally significant after Holm-correction (difficult: t_44_ = 2.200, p_Holm_ = 0.079, d = 0.176; impossible: t_44_ = 2.301, p_Holm_ = 0.079, d = 0.184), but significant when uncorrected (difficult: t_22_ = 2.309, p = 0.031, d = 0.481; impossible: t_22_ = 2.310, p = 0.031, d = 0.482). These results, together with those from Experiments 1 and 2, suggest that eye movements are influenced differently by listening challenges than pupil size, and that eye movements are unlikely driven by acoustic factors.

## Discussion

Pupillometry is commonly used to assess effort, but it has disadvantages. Here, we investigated the degree to which a new measure of effort – that is, the dispersion of eye movements – reflects effort across three listening experiments. Younger and older adults listened to speech masked by background babble at easy, difficult, and impossible speech-comprehension levels. We expected a u-shaped pattern in pupil size and gaze dispersion indicative of high effort for difficult comprehension, but low for easy and impossible speech comprehension. In the latter case, we predicted that individuals would give up investing effort when comprehension was impossible. The pupil-size data showed the expected u-shaped profile, whereas gaze dispersion decreased with increasing speech masking. The decrease in gaze dispersion for impossible speech comprehension was independent of stimulus acoustics, thus suggesting that gaze dispersion is governed by a different neuro-cognitive mechanism than pupil size.

### Pupil size reflects effort

Measurements of pupil size are frequently used to assess effort during task engagement, such as during listening to masked or degraded speech (Nicolai D. Ayasse & Wingfield, 2020; Kadem et al., 2020; Koelewijn, Zekveld, Lunner, & Kramer, 2018; Wendt, Dau, & Hjortkjær, 2016; Matthew B. Winn & Moore, 2018; M. B. Winn et al., 2018; Adriana A. Zekveld et al., 2014; Adriana A. Zekveld & Kramer, 2014; A. A. Zekveld et al., 2010). Our results are consistent with this previous work, showing that the pupil size is largest under difficult but manageable speech-comprehension conditions, whereas it is smaller when speech comprehension is easy or impossible (Ohlenforst et al., 2018; Ohlenforst et al., 2017; Wendt et al., 2018; Adriana A. Zekveld & Kramer, 2014). Although the current pupil-size results show the u-shaped profile, this profile appears a bit variable; for example, it was not significant for younger adults in Experiment 3 (note that pupil-size variability generally appeared larger in younger than older adults; error bars in Figures 2, 4, 5). The variability in the u-shaped profile of the pupil size may be partly related to the moving dot display that was presented during sound presentation to facilitate sensitivity to changes in eye movements. Pupillometry is typically recorded while individuals maintain fixation on a point on a computer monitor during speech listening (Farahani et al., 2020; Matthew B. Winn & Teece, 2021; Adriana A. Zekveld et al., 2014; Zhao et al., 2019), but this can alter speech processes related to mental imagery and comprehension (Cui & Herrmann, 2023; Johansson et al., 2012). Nonetheless, pupil size was associated with cognitive investment in the current study, reaffirming the utility of this marker in assessing listening effort.

### Eye movements reflect listening challenges differently than the pupil size

Recent works suggest that reductions in eye movements may index effort. Eye movements decrease when individuals perform a high-load compared to a low-load memory task (Hutton & Tegally, 2005; Kosch et al., 2018; Lipton, Frost, & Holzman, 1980; Walter & Bex, 2021) and when speech masking increases (Contadini-Wright et al., 2023; Cui & Herrmann, 2023). We also found that gaze dispersion decreased with decreasing SNR (Figures 2, 4, and 5), and extend this previous work by showing that the sensitivity of eye movements to changes in speech masking does not differ between younger and older adults. The absence of an age-group difference suggests that measuring gaze dispersion may be useful across different populations.

Previous studies that investigated changes in eye movements related to effort only included easy and difficult task conditions, but a signature of effort must also be sensitive to disengagement that occurs under conditions that exceed a person’s mental resources. We thus predicted a u-shaped effort profile when the range of easy to impossible listening conditions were used (Ohlenforst et al., 2018; Ohlenforst et al., 2017; Wendt et al., 2018; Yerkes & Dodson, 1908; Adriana A. Zekveld & Kramer, 2014). The results of the current study provide no indication of a u-shaped profile for gaze dispersion (Figures 2, 4, and 5). Instead, gaze dispersion decreased with decreasing SNR. On first glance, this may indicate that changes in eye movements are coupled to changes in the acoustic properties of the auditory environment rather than to effort per se. That is, sound intensity during sentence presentation increased with increasing SNR, because the babble was played at the same level across trials to which a sentence was added at different levels to create the three SNRs (cf. Cui & Herrmann, 2023; Kadem et al., 2020; Ohlenforst et al., 2017). Critically, acoustic differences ceased at sentence offset, after which only the babble continued. Gaze dispersion did not differ between easy and difficult speech-comprehension conditions after sentence offset, suggesting that, in the difficult condition, individuals stopped investing cognitively after a sentence ended. Yet, gaze dispersion during impossible speech comprehension remained reduced after sentence offset relative to easy and difficult comprehension. Moreover, in Experiment 3, in which participants could more readily anticipate the speech-comprehension challenge of the upcoming sentence, gaze dispersion for both the difficult and the impossible SNR were reduced prior to sentence onset. Acoustic differences thus do not appear to drive the reduction in the spatial extent of eye movements. Likewise, prior knowledge about the upcoming challenge in speech comprehension did not affect this pattern of results. The difference in the data patterns for pupil size and gaze dispersion may therefore reflect the underlying influence of distinct processes during challenging listening conditions.

### The role of different neurophysiological mechanisms in the changes in pupil size and eye movements

A wide network of brain regions underlies oculomotor functions, including the execution of eye movements and pupil dilation/constriction. Brain regions associated with eye movements include the superior colliculus, thalamus, basal ganglia, cerebellum, visual cortex, frontal and supplementary eye fields, anterior cingulate cortex, posterior parietal cortex, and the prefrontal cortex (Lencer & Trillenberg, 2008; Pierce, Clementz, & McDowell, 2019; Pierrot-Deseilligny, Milea, & Müri, 2004; Sparks, 2002). Pupil size is regulated by some of these same brain regions (Burlingham, Mirbagheri, & Heeger, 2022; Joshi & Gold, 2020; Wang, Boehnke, White, & Munoz, 2012; Wang & Munoz, 2021), but is also driven by regions that do not appear to play a prominent role in regulating eye movements, such as the locus coeruleus (Joshi & Gold, 2020; Mathôt, 2018; Strauch et al., 2022).

Critically, changes in pupil size have been linked to changes in arousal associated with locus coeruleus function (Bradley, Miccoli, Escrig, & Lang, 2008; Burlingham et al., 2022; Mathôt, 2018; Ross & Van Bockstaele, 2021; Wang et al., 2018). Here, too, the arousal system may underlie the observed pattern of pupil responses. In Experiment 1, in which the comprehension difficulty could not be predicted for a given trial, high SNR sentences elicited a transient pupil response (despite being highly intelligible; Figure 2). This transient pupil response was less prominent in Experiment 2 when individuals could predict, and thus were less surprised by, the SNR of a sentence (Figure 4). Moreover, there was a transient increase in pupil size following the visual cue that indexed the impossible SNR condition, possibly reflecting surprisal which, in turn, elicited arousal (Figure 4).

Eye movements can be automatically triggered or volitionally initiated (Pierce et al., 2019). Eye-movement reductions with increasing task load may include involuntary eye movements, as suggested by observations of task-difficulty related eye-movement reductions during fixation on a point on a computer monitor (Contadini-Wright et al., 2023; Cui & Herrmann, 2023). However, individuals may also voluntarily reduce eye movements under high cognitive demands, for example, to free resources associated with eye movements, avoid processing irrelevant information, and direct attention internally during listening (stereo listening via headphones is devoid of localizable external sound sources). A voluntary reduction in eye movement explorations may involve cognitive control and monitoring regions, such as the prefrontal, cingulate, and parietal cortices (Todd S. Braver, 2012; Cole & Schneider, 2007; Niendam et al., 2012), that drive structures associated with eye movement initiation, such as the frontal and supplementary eye fields and the superior colliculus (Pierce et al., 2019).

The pattern of gaze dispersion for the difficult SNR is consistent with the idea that eye movements are responsive to changes in effort. Gaze dispersion was reduced in the difficult condition, relative to the easy condition (Contadini-Wright et al., 2023; Cui & Herrmann, 2023), and then increased to similar levels as in the easy condition after sentence offset (Figures 2, 3, and 5). Yet, the continued reduction in gaze dispersion for the impossible condition suggests that eye movements may also be responsive to other aspects of cognition, such as monitoring. Participants could only listen to the babble noise during impossible trials and may have been trying to monitor the babble to discern speech patterns without necessarily investing effort. Effort is perhaps only invested after a listener successfully discerned speech patterns, when trying to suppress the babble to comprehend speech. Participants could not hear when the sentences ended in the impossible condition, likely leading them to continue to reduce eye movements throughout the trial. We would predict that in a study in which the babble noise continued for longer following sentence offset, visual exploration in the impossible condition would eventually increase and be similar to that of the easy condition. All together, we suggest that changes in the spatial dispersion of eye movements may be due to the internal direction of attention, and the avoidance of external visual information, irrespective of effort investment, whereas changes in pupil size indicate effort more directly through the involvement of arousal systems.

## Conclusions

The current study investigated the degree to which changes in eye movements reflect mental effort associated with listening in younger and older adults. Individuals listened to speech masked by background babble at signal-to-noise ratios associated with easy, difficult, and impossible speech comprehension. We expected individuals would invest little effort for easy and impossible speech (‘giving up’) but that they would exert higher effort for difficult speech. Consistent with this effort profile, a u-shaped pattern was observed for the pupil size, such that pupil size was largest for difficult comprehension, but lower for easy and impossible comprehension. In contrast, gaze dispersion decreased with increasing speech masking. However, gaze dispersion during difficult comprehension returned to levels similar to easy comprehension after sentence offset, during which acoustic stimulation was similar across conditions, whereas the spatial dispersion of eye movements during impossible comprehension continued to decrease. These findings suggest that acoustic factors do not drive changes in eye movement explorations. Rather, our data instead suggest that a cognitive process, different from the arousal-related systems regulating the pupil size, is involved in reduced eye movement explorations during high cognitive load. The current study highlights how effort in one sensory modality (audition) impacts the functional properties in another sensory modality (vision).

## Data availability

Data will be available upon reasonable request to the corresponding author.

## Author Contributions

BH: Conceptualization, methodology, validation, formal analysis, investigation, data curation, writing - original draft, writing - review & editing, visualization, supervision, project management, funding acquisition. JDR: Conceptualization, methodology, investigation, writing - review & editing.

## Acknowledgements

The research was supported by the Canada Research Chair program (CRC-2019-00156, 232733), the Natural Sciences and Engineering Research Council of Canada (NSERC Discovery Grant: RGPIN-2021-02602), the Canadian Institutes of Health Research (CIHR: 186236), and the William Demant Foundation. We thank Tazeen Atif and Priya Pandey for help during data collection. We thank Ingrid Johnsrude and Lauren Fink for fruitful discussions.

## Statements and Declarations

The author has no conflicts or competing interests.

